# Both HIV and Tat Expression Decrease Prepulse Inhibition with Further Impairment by Methamphetamine

**DOI:** 10.1101/2020.06.11.130666

**Authors:** T. Jordan Walter, Jared W. Young, Morgane Milienne-Petiot, D. S. Deben, Robert K. Heaton, Scott Letendre, David J. Grelotti, William Perry, Igor Grant, Arpi Minassian, Translational Methamphetamine AIDS Research Center (TMARC)

## Abstract

HIV infection and methamphetamine (METH) use are highly comorbid and represent a significant public health problem. Both conditions are known to negatively impact a variety of brain functions. One brain function that may be affected by HIV and METH use is sensorimotor gating, an automatic, pre-conscious filtering of sensory information that is thought to contribute to higher order cognitive processes. Sensorimotor gating is often measured using prepulse inhibition (PPI), a paradigm that can be conducted in both humans and animals, thereby enabling cross-species translational studies. While previous studies suggest HIV and METH may individually impair PPI, little research has been conducted on the effects of combined HIV and METH on PPI. The goal of this cross-species study was to determine the effects of METH on PPI in the inducible Tat (iTat) mouse model of HIV and in people with HIV. PPI was measured in the iTat mouse model before, during, and after chronic METH treatment and after Tat induction. Chronic METH treatment decreased PPI in male but not female mice. PPI normalized with cessation of METH. Inducing Tat expression decreased PPI in male but not in female mice. No interactions between chronic METH treatment and Tat expression were observed in mice. In humans, HIV was associated with decreased PPI in both men and women. Furthermore, PPI was lowest in people with HIV who also had a history of METH dependence. Overall, these results suggest HIV and METH may additively impair early information processing in humans, potentially affecting downstream cognitive function.

**HIGHLIGHTS:**

- HIV decreased PPI in men and women
- PPI was most decreased in people with HIV and a history of METH dependence
- Chronic METH treatment decreased PPI in male but not female mice
- Tat expression decreased PPI in male but not female mice
- Chronic METH treatment and Tat expression did not interact to affect PPI in mice

## 1. INTRODUCTION

Methamphetamine (METH) use is highly prevalent among people with HIV and represents a significant public health concern. Indeed, one study found that 31% of people in outpatient HIV clinics used METH in the past year (Clark et al., 2012), and another found that 13% of HIV+ adults have a METH use disorder (Hartzler et al., 2017). Both HIV (Walker and Brown, 2018) and METH use (Potvin et al., 2018) negatively impact a variety of brain functions. One brain function that may be impaired by combined HIV and METH use is sensorimotor gating, a normal process of sensory information filtering. Sensorimotor gating is an involuntary, pre-conscious process that is thought to impact downstream cognitive functions (Geyer, 2006). Sensorimotor gating has been included by the National Institute of Mental Health (NIMH) in its Research Domain Criteria (RDoc) because of its relevance to mental health and psychopathology (Harrison et al., 2019). Deficits in sensory processing of internal and/or external stimuli may contribute to symptoms as diverse as cognitive deficits, affective dysregulation, somatization, and visual/auditory hallucinations (Harrison et al., 2019). Indeed, decreased sensorimotor gating has been observed in several neuropsychiatric conditions, including schizophrenia (Braff and Geyer, 1990; Minassian et al., 2007) and bipolar disorder (Perry et al., 2001).

Sensorimotor gating is often measured using prepulse inhibition (PPI) paradigms, in which a weak stimulus (a prepulse) is presented shortly before a larger, startle-inducing stimulus (a pulse), thereby reducing the magnitude of the startle response (Braff et al., 2001). Importantly, PPI can be measured in animals as well as humans, thus enabling translational studies to determine mechanisms underlying PPI deficits and develop treatment strategies. Overall, PPI is an objective, quantitative, and translational measure of pre-conscious information processing that may inform our understanding of downstream cognitive processes in populations such as HIV+ people who use METH.

While previous studies have examined the individual effects of HIV and METH on PPI, little is known about how combined HIV and METH affects PPI. Further understanding the effects of HIV and METH on CNS processes such as sensorimotor gating will provide insight into mechanisms of psychopathology and potential treatment strategies. A previous study found PPI was not significantly different in people with HIV compared to controls (Minassian et al., 2013); however, a preclinical study found PPI was decreased in female mice of the gp120 mouse model of HIV (Henry et al., 2014). While there are no known studies examining the effects of METH on PPI in humans, preclinical studies find that acute and chronic METH intoxication decreases PPI (Arai et al., 2008; Chao et al., 2012). Alternatively, withdrawal from chronic METH treatment increases PPI (Henry et al., 2014). One preclinical study examined the effects of METH withdrawal in the gp120 mouse model of HIV and found no interaction between gp120 expression and METH withdrawal (Henry et al., 2014). Studies in additional animal models of HIV would provide useful information, as the gp120 model is limited by the constitutive expression of only one of the HIV proteins. Another such animal model of HIV is the inducible Tat (iTat) mouse, which expresses the HIV protein Tat in the brain, thereby replicating many of the neurotoxic effects of HIV seen in humans (Langford et al., 2018). One advantage of using the iTat model in studies of HIV is that Tat expression is induced only after treatment with doxycycline, thereby allowing the researcher to model acquired HIV pathology in adulthood, and avoiding the potential conflicting effects of the presence of HIV proteins during development. The iTat mouse model can be studied in parallel with HIV+ humans to determine the effects of quantifiable levels of METH on PPI in HIV.

Given the prevalence of METH dependence in people with HIV and the unknown but potentially interactive effects of HIV and METH on sensorimotor gating, the objective of this study was to determine the effects of HIV and chronic METH on PPI using a translational, cross-species approach. We performed animal studies using the iTat mouse model of HIV to evaluate if these iTat mice, after induction with doxycylcline, would exhibit greater PPI deficits relative to their wildtype littermates and if previous chronic METH treatment would exacerbate these deficits. We also studied human participants with HIV and a history of METH dependence and hypothesized these conditions would additively or synergistically decrease PPI. We also assessed characteristics of HIV (e.g., CD4+ counts, viral load) and METH use (e.g., total days of METH use, quantity of METH use) to determine whether more severe illness and drug use history features were associated with poorer sensorimotor gating. Since PPI is thought to contribute to higher order cognitive function, we also examined neuropsychological function across several domains (e.g., attention, memory) and hypothesized that worse neuropsychological function would be correlated with worse PPI. Such findings would provide valuable insight into mechanisms of sensorimotor gating deficits and enable the testing of possible therapeutic options.

## 2. MATERIALS AND METHODS

The human and animal components of this study were a part of the Translational Methamphetamine AIDS Research Center (TMARC), a multi-project program of research focused on understanding the combined effects of HIV and METH dependence on brain structure and function. The UCSD Institutional Review Board and Human Research Protections Program approved the human components of the study. Subjects gave written informed consent for inclusion in the trial. Animal components were approved by the UCSD Institutional Animal Care and Use Committee and conformed to NIH guidelines.

### 2.1 Animal Subjects

The mouse experiments were part of a larger examination of the individual and combined effects of HIV-1 and METH conducted by the TMARC. In this study 120 mice were used, 60 males and 60 females. All animals were housed with a maximum of four mice per cage and maintained in a temperature-controlled vivarium (21±1 °C) on a reversed day-night cycle (lights on at 7:00 PM, off at 7:00 AM). All mice had *ad libitum* access to water and were food-restricted to approx. 85% of their free-feeding body weight. Testing occurred during the dark phase of the day-night cycle between 1:00 PM and 6:00 PM. All mice were genetically modified with the TGFAP+ promotor and only half of them also had the inducible Tat86+ promotor. All mice were in quarantine for 8 weeks before they were brought to the lab. The males and females were tested on separate days.

### 2.2 Animal Drug Treatment

METH injections were obtained by dissolving methamphetamine hydrochloride (Sigma, St Louis, MO) in saline (0.9%) and were administered subcutaneously at 5 ml/kg injection volume. 25-day regimen was with an escalating dose, starting from 1mg/kg at day one up to 6 mg/kg at day 25 (Fig 1A,B). During the regimen, mice were weighed twice a week to determine the injection volume for the following days. Thus, the aim of this study was to investigate the effects of long term METH exposure rather than the acute high-dose effects, thereby modeling the human condition of METH dependence. Similar doses and injection volumes were used for the control (saline) condition.

**Figure 1.**
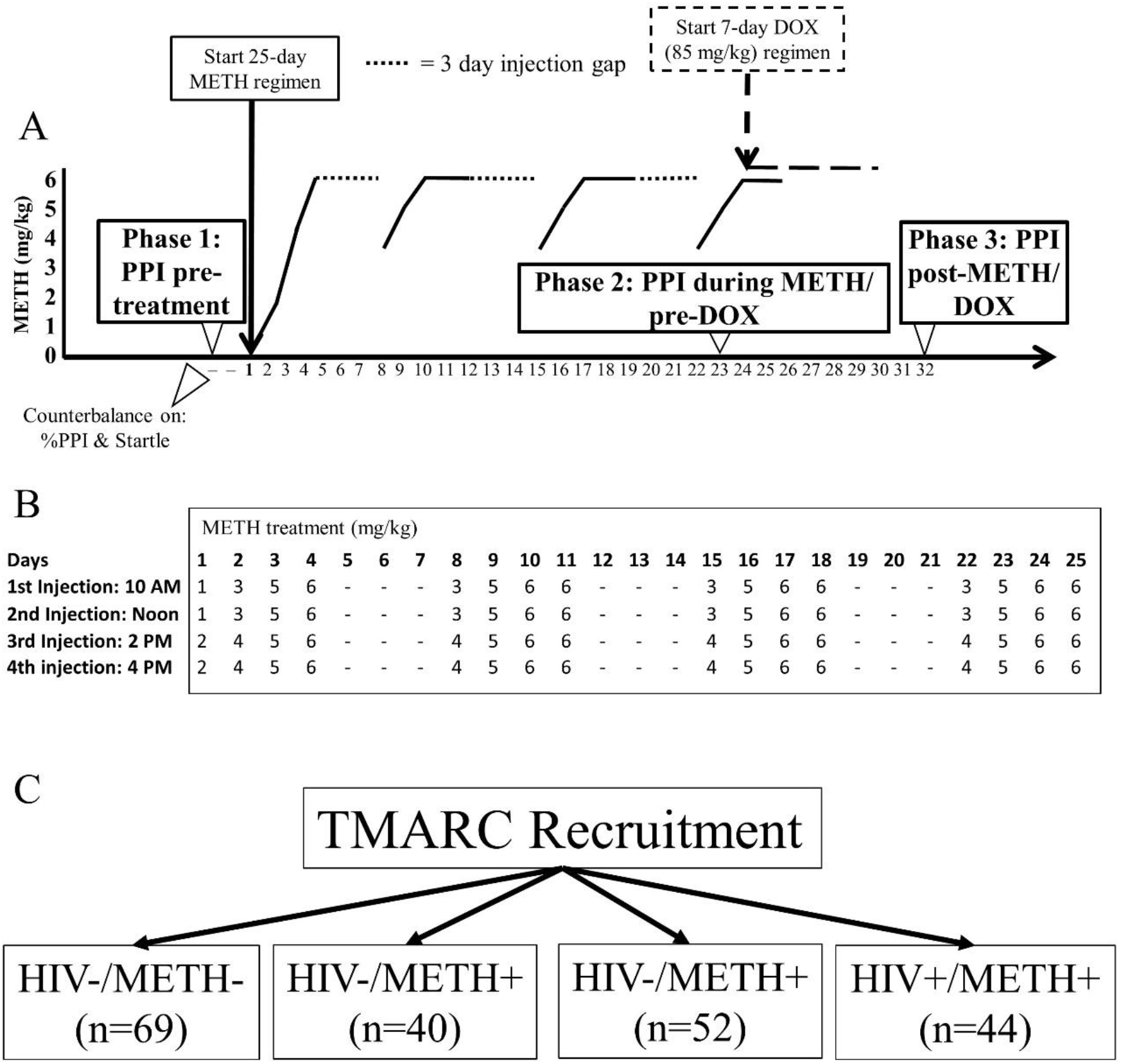
Schematic of experimental design. A.) WT and iTat mice were assessed for PPI at baseline (Phase 1: pre-METH and pre-DOX) and matched into four groups (WT+saline, WT+METH, Tat+saline, Tat+METH; n=15/group) based on percent PPI and startle reactivity. Two days later, mice began a 25 day regimen of injections with either saline or METH. Mice were assessed again for PPI on the 23^rd^ day of METH treatment (Phase 2: during METH and pre-DOX). The next day, a 7 day DOX treatment (85 mg/kg) was started. Mice were assessed for PPI a final time (Phase 3: post-METH and DOX) two days after completion of the DOX regimen. B.) METH was administered four times per day in an escalating manner, starting with 1 mg/kg and ending at 6 mg/kg. Injections were given for four days, followed by three days of no injections. This pattern was repeated a total of four times. C.) Participants with or without HIV and with or without a history of METH dependence were recruited for the human studies.

Doxycycline (DOX) injections were obtained by dissolving doxycycline hydrochloride (Sigma, St Louis, MO) in saline, given at a dose of 100 mg/kg and administered intraperitoneally. Injections were given in a volume of 10 ml/kg body weight. The doxycycline regimen started on the second to last day of the METH regimen. DOX injections were given for 7 consecutive days once daily to induce the Tat protein. This regimen was based on DOX dose-effect relationships on Tat expression as investigated in previous studies (Kesby et al., 2018; Kesby et al., 2017). Fresh syringes were used for every injection given to each mouse and were prepared with a maximum of two hours before the injections were given. Animal experimenters were blinded to the genotype, but not METH treatment of animals. This was because these same individuals made up the METH solutions for injection and because of the obvious behavioral effects of the METH.

### 2.3 Animal Study Outline

Power analyses showed a minimum of 12 mice per group were required for an ANOVA to have 96% power to detect drug and genotype effects based previous studies at an α=0.05 level. The animal studies consisted of 3 phases (Fig 1A,B). Phase 1 was testing at baseline, before any METH or DOX treatments to determine whether any differences between genotypes existed prior to Tat activation. Phase 1 PPI testing occurred 2 days before METH treatment began. Next, mice were matched into four test groups of 15 mice each (1=WT+saline 2=WT+METH, 3=Tat+saline, 4=Tat+METH) based on startle reactivity, per our standard PPI methodology for rodents (Henry et al., 2014). Phase 2 started after matching. During Phase 2, half of the mice received METH injections for 25 days, 4 injections a day, the other half were injected in the same manner with saline. Phase 2 PPI testing occurred on the 23^rd^ day of METH administration, one day before DOX administration began. Phase 3 started after Phase 2, with a 2-day overlap between the two phases. During Phase 3, mice received DOX injections for seven days to express the Tat-protein. The final PPI test was done two days after completion of the DOX injections. The entirety of the animal experiment was performed once.

### 2.4 Animal Prepulse Inhibition

PPI was measured in mice as described in detail previously (Henry et al., 2014; Young et al., 2014). The mice acclimated to the testing room for at least 30 minutes before they were placed into the testing chambers. The experiment consisted of a 5 min acclimatization period inside the PPI chamber with a background noise of 65-dB, followed by a 15 min test session of PPI. Five different trials were performed, including a 40-ms, 120-dB startle pulse (P120), three 20-ms prepulse + P120 pulse combinations with different intensities of the prepulses (68, 71 or 77 dB) and no-stimulus trials, composed of accelerometer recordings achieved in the absence of any stimulus and served to assess baseline motor activity. The interval was 100 ms between prepulse and pulse. The inter trial interval (ITI) had an average duration of 15 s, varying between 7 and 23 s. Between each of these trials, no-stimulus trials were interchanged. Additionally, six P120 trials were introduced at both the beginning (Block 1: to assess startle reactivity before appreciable habituation) and the end of the acoustic test session (Block 2: to assess habituation across the session by comparison with Block 1). For each trial type presentation, the mean startle magnitude was determined by averaging 100 ms readings taken from the onset of the P120 stimulus.

Hence, the groups of assessments within the PPI tests included: 1.) PP100s, which included a prepulse of 69, 73, and 81, to a pulse dB of 100, 2.) PP120s, which included a prepulse of 69, 73, and 81, to a pulse dB of 120, 3.) pulse intensity block, which included random pulses at dB of 69, 73, 81, 90, 100, 110, and 110, 4.) ISI curves, which included a prepulse of 73 dB and a pulse dB of 120, with the inter-stimulus interval (ISI) varied for 25, 50, 100, 200, or 500 ms between prepulse and pulse, and 5.) a habituation (HAB) curve to repeated 120 dB pulses. See the Supplemental Material for statistics on PP100s, pulse intensities, ISI curves, and habituations curves.

### 2.5 Human Participants

Four groups of participants were examined: HIV−/METH− (n = 69), HIV−/METH+ (n = 40), HIV+/METH− (n = 52), and HIV+/METH+ (n = 44) (Fig 1C). Recruitment goals were n=80/group. This design afforded 80% power to detect a main effect of HIV or METH of Cohen’s d ≥ 0.363 at the α=0.05 level and an interaction of Cohen’s d ≥ 0.638 or larger at the α=0.05 level. While these recruitment goals were reached for the parent TMARC study, not all participants underwent PPI testing. HIV status was determined by enzyme-linked immunosorbent assay (ELISA) and a confirmatory Western blot. A history of METH use disorder was determined via the DSM-IV criteria for METH abuse and dependence as assessed by the Composite International Diagnostic Interview Version 2.1. The METH+ groups included participants who met criteria for both lifetime METH dependence and METH abuse or dependence within the past 18 months.

Potential participants were excluded if they reported histories of psychosis or significant medical conditions, such as hepatitis C infection, or neurological conditions, such as seizure, stroke, or multiple sclerosis, that are known to affect cognitive functions. Participants were also excluded if they met diagnostic criteria for current abuse or dependence on alcohol or any illicit drugs. Participants who tested positive for METH or any other illicit drugs (aside from cannabis) were excluded. The parent study from which this investigation was derived (TMARC) does not exclude subjects with positive cannabis tests because some antiretroviral medications produce false positives on cannabis screening tests and, since cannabis can be detected in urine for up to one month, positive screens do not necessarily indicate recent use.

### 2.6 Human Prepulse Inhibition Procedures

Standard procedures for PPI were implemented as described previously in detail (Feifel et al., 2009; Minassian et al., 2007; Minassian et al., 2013; Perry et al., 2004; Perry et al., 2001; Perry et al., 2007). Participants refrained from nicotine and caffeine use 30 min before startle testing and underwent a brief hearing screening to ensure that they could hear 500, 1000, and 6000 Hz tones bilaterally. During the procedure, participants were seated in a reclining chair. The eyeblink component of the auditory startle reflex was measured using electromyography (EMG) of the orbicularis oculi muscle as previously described (Perry et al., 2001). Two miniature electrodes (InVivo Metric, Healdsburg, CA) were positioned below and to the right of the subject’s right eye, and a ground electrode was placed behind the right ear over the mastoid. Subjects were instructed to keep their eyes open and fixed on a square on the wall. Acoustic startle and prepulse stimuli were presented through headphones (Model TDH-39-P, Maico, Minneapolis, MN) and electromyographic activity recorded by the electrodes was directed through a customized EMG amplifier to a computerized startle response monitoring system for digitization and analysis (SR-LAB, San Diego Instruments, San Diego, CA.).

The startle session was conducted as previously described (Minassian et al., 2013). It began with a 5-min acclimation period of 70 dB white noise followed by four blocks of trials. The first and last blocks consisted of five pulse-alone trials of 40 ms 115-dB startle stimuli. Blocks two and three consisted of three pulse-alone and nine prepulse-pulse trials per block presented in pseudorandom order. The 20-ms prepulse stimuli preceded the startle stimulus by 60 ms (onset-to-onset) and were either 74, 78, or 86 dB (i.e. – 4, 8, or 16 dB above the 70 dB background noise). The inter-trial interval averaged 15 s with a range of 11 to 21 s. All blocks contained hidden “no stimulus” trials where no sound was delivered but EMG data were collected. The session duration was approximately 15 min. The experimenter was not blinded to the group of the participants.

### 2.7 Neuropsychological Testing

The neurocognitive battery included tests designed to assess verbal fluency (Letter Fluency, Animal Fluency, and Verb Fluency), executive functioning (Trail Making Test Part B and the Wisconsin Card Sorting Test-64), speed of information processing (WAIS-III Digit Symbol and Symbol Search subtests, Stroop Color test, and Trail Making Test Part A), learning (Hopkins Verbal Learning Test-Revised (HVLT-R) and Brief Visual Memory Test-Revised (BVMT-R), memory (delayed recall of HVLT-R and BVMT-R), working memory (Paced Auditory Serial Addition Test-50 and WMS-III Spatial Span), and complex motor skills (Grooved Pegboard). Raw scores were converted to T-scores using published, demographically adjusted normative standards. T-scores were then converted to deficit scores, which range from 0 (T > 39; no impairment) to 5 (T < 20; severe impairment), with higher deficit scores reflected greater neurocognitive disturbance. Deficit scores from primary test measures were then averaged to derive the Global Deficit Score (GDS) (Carey et al., 2004).

### 2.8 Data Processing and Statistical Analyses

For the animal experiments, we calculated the percentage decrease in startle response in the three different prepulse + pulse trials compared to the pulse trial without prepulse. The following formula was used for calculation: ([1 − (startle magnitude on prepulse + pulse trials/startle magnitude on P120 trials)] × 100). In addition, the startle reactivity and startle habituation were calculated. The average startle magnitude over the record window (i.e., 65 msec) was used for all data analysis. PPI was assessed using mixed analysis of variance (ANOVA) with METH and genotype as between-subject factors. The data were analyzed for outliers and data points two standard deviations beyond the mean were removed from analysis. Data from each testing block were analyzed separately, and the data were split by sex given the different testing times. Each time period (baseline, during METH, and after Tat expression), was analyzed separately. Follow-up ANOVAs were conducted on significant effects and interactions. Data were analyzed using BMDP Statistical software (Statistical Solutions Inc., Saugus, MA), with significant difference set at p<0.05.

For the human experiments, PPI was calculated as the percent decrement in startle amplitude in the presence of the prepulse compared to the amplitude without the prepulse [100 – (prepulse amplitude/pulse amplitude × 100)]. PPI was calculated for each of the three prepulse conditions in the blocks containing prepulse trials, and average PPI was calculated by averaging the PPI for each of the three prepulse conditions in those blocks. Subjects with an average amplitude to the pulse alone trials that was less than three times the average amplitude for no-stimulus trials in any of the four blocks were classified as startle non-responders and were excluded from further analyses. Subjects with a greater startle response to the prepulse trials than the pulse alone trials were classified as prepulse facilitators and their negative PPI values were set to zero.

Statistical analyses for the human studies were carried out using the Statistical Package for the Social Sciences (SPSS) 26. The PPI data were inspected for outliers. Outliers were defined as values less than 1.5 interquartile intervals below the 1^st^ quartile (25^th^ percentile) or 1.5 interquartile intervals above the 3^rd^ quartile (75^th^ percentile) for each group. This was done according to a commonly used definition of outliers (Tukey, 1977). A total of seven outlying data point were identified using this method and were excluded from subsequent analysis. Outliers were only assessed for the men, as there were already few women participants. PPI was assessed using a mixed analysis of variance (ANOVA) with prepulse intensity as within-subjects factors and HIV status and history of METH dependence as between-subjects factors. We also used a one-way ANOVA with Dunnett’s posthoc test comparing to the HIV−/METH− group to examine statistical differences between average PPI across the four groups. METH use characteristics, HIV illness features, and neuropsychological data were non-normally distributed, therefore the non-parametric Spearman’s rho was used to calculate correlations between these variables and percent PPI for each prepulse intensity. Statistical significance was defined as p<0.05.

## 3. RESULTS

Data from male and female mice were analyzed separately because the experiments for male and female mice were conducted separately.

### 3.1 PPI Analyses in Male Mice – Phase 1: Baseline pre-METH & pre-DOX

We observed a main effect of prepulse intensity on PPI [F(2,116)=174.4, *p*<0.0001] (Fig 2A). PPI at each prepulse intensity level was significantly higher than the previous level (PPI at 69<73<81). There was no effect of genotype, or genotype-by-prepulse intensity interaction on PPI (F<1.8, ns).

**Figure 2.**
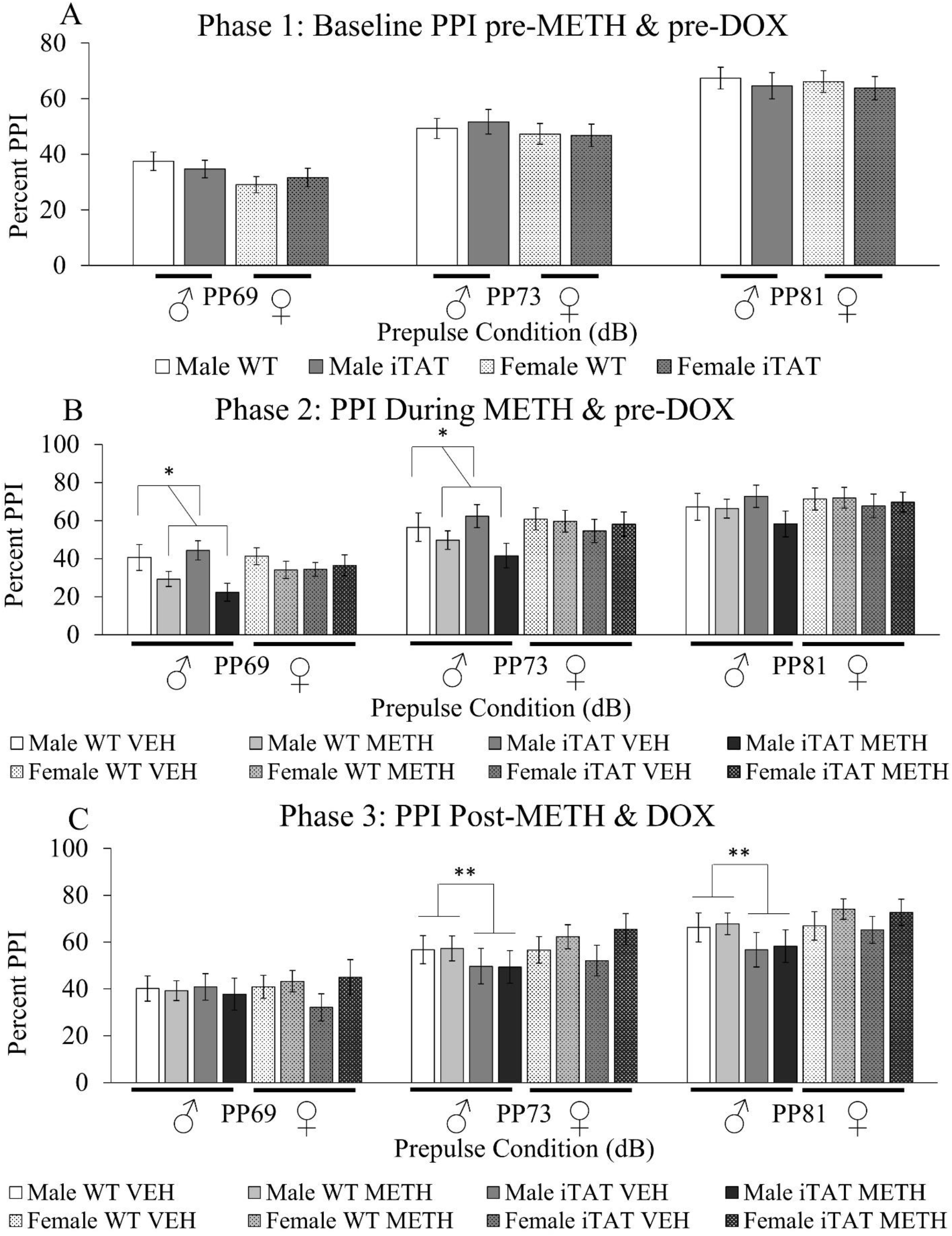
PPI testing in mice pre-METH and pre-DOX, during METH and pre-DOX, and post-METH and post-DOX. A.) WT and iTat male and female mice (n=30/group) were assessed for PPI prior to METH or DOX treatment. B.) Mice then received a 25-day regimen of either saline or METH injections and were assessed for PPI on the 23^rd^ day of METH treatment. C.) Mice then received a 7-day course of DOX injections overlapping by two days with the 25 day METH treatment. Two days after the last DOX injection, mice were assessed for PPI. Data are presented as mean ± S.E.M. * = main effect of METH (ANOVA, p<0.05). ** = main effect of Tat induction (ANOVA, p<0.05)

### 3.2 PPI Analyses in Male Mice – Phase 2: During METH & pre-DOX

A main effect of prepulse intensity on PPI [F(2,110)=171.0, *p*<0.0001] (Fig 2B) was again observed. PPI at each prepulse intensity level was significantly higher than the previous level (PPI at 69<73<81). A main effect of METH on PPI was observed [F(1,55)=5.6, *p*<0.05], revealing that those on METH exhibited lower PPI irrespective of genotype (Fig 2B). No effect of genotype, genotype-by-prepulse intensity, or genotype-by-prepulse intensity-by-METH interaction on PPI was observed (F<1, ns).

### 3.3 PPI Analyses in Male Mice – Phase 3: After METH & DOX

There was a main effect of prepulse intensity on PPI [F(2,94)=96.6, *p*<0.0001] (Fig 2C). PPI at each prepulse intensity level was significantly higher than the previous level (PPI at 69<73<81). METH administration had been stopped at this point, and no main effect of genotype, METH, METH-by-prepulse intensity, or METH-by-genotype-by-prepulse intensity on PPI was observed (F<1, ns). Interestingly, a genotype-by-prepulse intensity interaction was observed [(F2,94)=4.2, *p*<0.05], revealing lower PPI in the Tat mice vs. WT littermates, specifically at higher prepulse intensities (73 and 81) (Fig 2C).

In summary, in male mice, during METH treatment, PPI was significantly lowered irrespective of genotype. After cessation of METH and administration of DOX, the METH effect disappeared but DOX resulted in impaired PPI in the iTat mice relative to their WT littermates at higher prepulse levels.

### 3.4 PPI Analyses in Female Mice – Phase 1: Baseline pre-METH & pre-DOX

A main effect of prepulse on PPI [F(2,114)=153.2, *p*<0.0001] (Fig 2A) was observed. PPI at each prepulse level was significantly higher than the previous level (PPI at 69<73<81). There was no effect of genotype, or genotype-by-prepulse interaction on PPI (F<1, ns).

### 3.5 PPI Analyses in Female Mice – Phase 2: During METH & pre-DOX

We observed a main effect of prepulse on PPI [F(2,110)=171.0, *p*<0.0001] (Fig 2B). PPI at each prepulse level was significantly higher than the previous level (PPI at 69<73<81). Unlike in the males, no main effect of drug was observed (F<1, ns). No effect of genotype, genotype-by-prepulse, or genotype-by-prepulse-by-METH interaction on PPI was observed (F<1, ns).

### 3.6 PPI Analyses in Female Mice – Phase 3: After METH & DOX

A main effect of prepulse on PPI [F(2,108)=131.3, *p*<0.0001] (Fig 2C) was again observed. PPI at each prepulse level was significantly higher than the previous level (PPI at 69<73<81). METH administration had been stopped, and no main effect of METH, genotype, METH-by-prepulse, or METH-by-genotype-by-prepulse on PPI was observed (F<2, ns). Unlike in the males, no genotype-by-prepulse interaction was observed (F<1, ns).

In summary, METH had no effect on PPI in females, nor did females exhibit the same Tat-driven drop in PPI after DOX treatment that was seen in the male mice.

### 3.7 Human Demographics

Because sex differences were observed in the animal studies, we analyzed the data from men and women separately. A total of 160 men were recruited: 42 in the HIV−/METH− group, 29 in the HIV−/METH+ group, 47 in the HIV+/METH− group, and 42 in the HIV+/METH+ group (Table 1).

**Table 1:** Demographic and Illness Features: Male Participants.

|  | **HIV-/**<br>**METH-**<br>**(n=42)** | **HIV-/**<br>**METH+**<br>**(n=29)** | **HIV+/**<br>**METH-**<br>**(n=47)** | **HIV+/**<br>**METH+**<br>**(n=42)** | **Statistics** |
| --- | --- | --- | --- | --- | --- |
| **Age (yrs)** | 35.7 ±<br>13.0<br>(18-64) | 37.0 ±<br>11.0<br>(21-60) | 41.0 ±<br>10.8<br>(23-70) | 39.4 ± 6.6<br>(24-54) | **HIV+ groups**<br>**> HIV-**<br>**groups** |
| **Ethnicity (n)** | 6 Afr.<br>Am.<br>10 Hisp.<br>0 Asian<br>25 Cauc.<br>1 Other | 8 Afr.<br>Am.<br>5 Hisp.<br>0 Asian<br>16 Cauc.<br>0 Other | 9 Afr.<br>Am.<br>13 Hisp.<br>2 Asian<br>22 Cauc.<br>1 Other | 4 Afr.<br>Am.<br>10 Hisp.<br>0 Asian<br>27 Cauc.<br>1 Other | NS |
| **Education (yrs)** | 14.4 ±<br>2.2<sup>A</sup><br>(10-20) | 12.7 ±<br>2.2<sup>B</sup><br>(9-18) | 14.2 ±<br>2.1<sup>A</sup><br>(9-18) | 14.1 ±<br>2.6<sup>A,B</sup><br>(8-20) | **Groups with**<br>**the same**<br>**letters are not**<br>**significantly**<br>**different** |
| **WRAT-4 Score** | 105 ± 11<sup>A</sup><br>(82-131) | 97 ± 9<sup>B</sup><br>(83-116) | 103 ±<br>11<sup>A,B</sup><br>(83-130) | 106 ± 13<sup>A</sup><br>(87-131) | **Groups with**<br>**the same**<br>**letters are not**<br>**significantly**<br>**different** |
| **Global Deficit Score** | 0.20 ±<br>0.21<br>(0-0.95) | 0.37 ±<br>0.39<br>(0-1.44) | 0.34 ±<br>0.30<br>(0-1.11) | 0.48 ±<br>0.45<br>(0-1.67) | **HIV+ groups**<br>**> HIV-**<br>**groups;**<br>**METH+**<br>**groups >**<br>**METH-**<br>**groups** |
| **BDI-II Total Score** | 2.3 ± 3.3<br>(0-13) | 13.7 ±<br>10.9<br>(0-35) | 11.6 ±<br>13.3<br>(0-58) | 13.7 ±<br>12.0<br>(0-47) | **HIV+ groups**<br>**> HIV-**<br>**groups;**<br>**METH+**<br>**groups >**<br>**METH-**<br>**groups** |
| **Lifetime Substance Use**<br>**Disorder (n)** | 20 | 29 | 22 | 42 | **Chi-squared:**<br>**p<0.0005** |
| **Age First METH use (yrs)** | - | 22.4 ± 11.0<br>(8-56) | - | 25.1 ± 7.6<br>(13-48) | NS |
| **Days since last METH use (days)** | - | 197 ± 407<br>(4-2191) | - | 117 ± 126<br>(1-426) | NS |
| **Total days METH use (days)** | - | 2262 ± 1961<br>(86-8797) | - | 1744 ± 1819<br>(59-9771) | NS |
| **Total quantity METH use (grams)** | - | 4131 ± 8182<br>(35-40185) | - | 1961 ± 4413<br>(12-26526) | NS |
| **Current CD4+ Cell Count** | - | - | 645 ± 301<br>(81 – 1712) | 563 ± 231<br>(82 – 1053) | NS |
| **Nadir CD4+ Cell Count** | - | - | 319 ± 220<br>(3-800) | 282 ± 182<br>(8-840) | NS |
| **Estimated Duration HIV Infection (yrs)** | - | - | 10.5 ± 9.2<br>(0.1-28.9) | 8.4 ± 6.4<br>(0.1-23.8) | NS |
| **Plasma Viral Load - log10(copies per mL)** | - | - | 2.11 ± 1.21<br>(0.00-5.45) | 2.39 ± 1.41<br>(0.00-5.95) | NS |
| **Current ART Use (n)** | - | - | 35 Yes<br>12 No | 31 Yes<br>11 No | NS |
| **Months Total ART Exposure** | - | - | 70.4 ± 78.2<br>(0-275) | 59.2 ± 58.0<br>(0-181) | NS |
| **Current Number of ARVs Using (n)** | - | - | 2.7 ± 1.7<br>(0-6) | 2.7 ± 1.7<br>(0-5) | NS |
| **Participants with AIDs (n)** | - | - | 18 | 17 | NS |

Alternatively, only 45 women were recruited: 27 in the HIV−/METH− group, 11 in the HIV−/METH+ group, 5 in the HIV+/METH− group, and 2 in the HIV+/METH+ group (Table S1). While the data for both sexes were analyzed, the data for the women are likely not generalizable given the very small number of participants in some of the groups. Analyses for women are therefore presented as supplementary material. Among men, there was a significant difference in age between groups, with HIV+ groups being older than HIV− groups. The groups were well-matched for ethnicity. The HIV−/METH+ group tended to have fewer years of education and lower WRAT-4 scores. The HIV+ groups and METH+ groups had higher Global Deficits Scores and higher Beck’s Depression Inventory II scores. The METH+ group also had a higher number of lifetime substance use disorders. The METH+ groups were well-matched for METH use characteristics and the HIV+ groups for HIV illness features.

### 3.8 PPI in Humans

In men, there was a main effect of prepulse intensity on average PPI [F(2,300)=194.49, p<0.0005] such that PPI increased with increasing prepulse intensity (Fig 3A).

**Figure 3.**
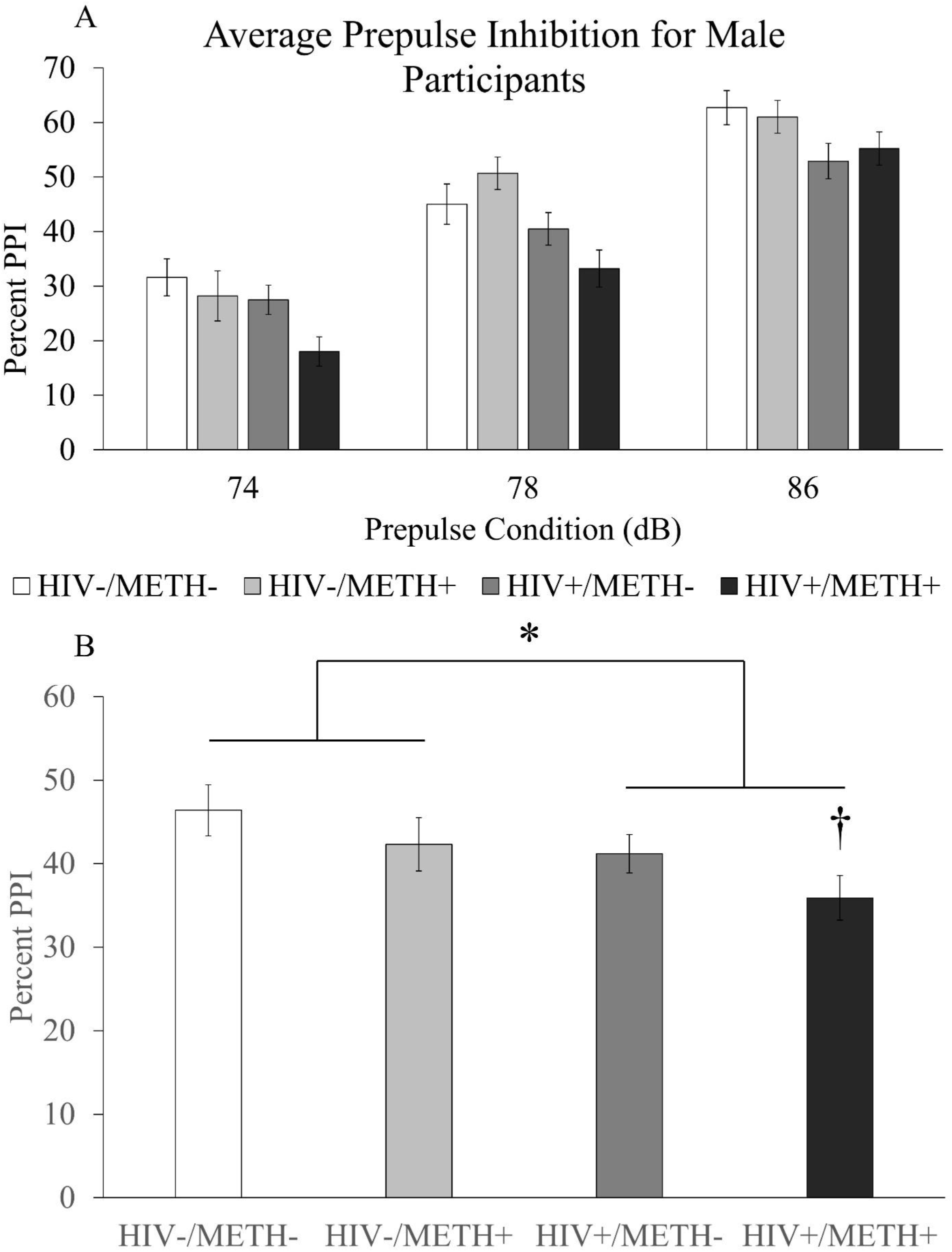
Effects of HIV and a history of METH dependence on PPI in male participants. Four groups of human men with or without HIV and with or without a history of METH dependence (HIV−/METH−: n=42, HIV−/METH+: n=29, HIV+/METH−: n=47, HIV+/METH+: n=42) were recruited and assessed for PPI at three different prepulse intensities. Average PPI for each group at each prepulse intensity are shown in panel A. PPI averaged across prepulse intensities are shown in panel B. Data are presented as mean ± S.E.M. * = main effect of HIV (Two-way ANOVA, p=0.003); † = significantly different compared to HIV−/METH− group (p<0.019, Dunnett’s posthoc test).

There was no interaction of prepulse intensity with HIV or prepulse intensity with METH; however, there was a significant prepulse-by-HIV-by-METH interaction [F(2,300)=3.59, p=0.029] (Fig 3A). There was also a main effect of HIV [F(1,150)=9.35, p=0.003] (Fig 3A,B), but no main effect of METH and no HIV-by-METH interaction. Mauchly’s test showed that the sphericity assumption was not violated (p=0.153). Because the ages of the HIV+ groups were significantly higher than the ages of the HIV− groups, we performed additional analyses with age as a covariate. In this model, there was still a main effect of prepulse intensity [F(2,298)=6.27, p=0.002] and a significant prepulse intensity-by-HIV-by-METH interaction [F(2,298)=3.80, p=0.023]. There was no prepulse intensity-by-HIV interaction or prepulse intensity-by-METH interaction. There was still a main effect of HIV [F(1,149)=8.08, p=0.005] and no effect of METH or HIV-by-METH interaction. Again, the sphericity assumption was not violated per Mauchly’s test (p=0.197).

Because there was no significant interaction between HIV and METH, we looked for evidence of an additive, detrimental effect of the two conditions on PPI. Parameter estimates for HIV status yielded a B value of 5.695 [F(1,156)=4.157, p=0.043, 95% confidence interval: 0.178 – 11.213], while the B value for METH use was 4.757 [F(1, 156)=2.900, p=0.091, 95% confidence interval: −0.761 – 10.275]. When age was included as a covariate, parameter estimates for HIV status yielded a B value of 6.121 [F(1,154)=3.578, p=0.060, 95% confidence interval: −2.274 – 14.517]. Parameter estimates for METH status yielded a B value of 5.378 [F(1,154)=2.749, p=0.099, 95% confidence interval: −2.032 – 12.789]. A one-way ANOVA between the four groups was nearly significant [F(3,155)=2.522, p=0.060]. Posthoc testing showed that PPI in the HIV+/METH+ group was significantly lower than PPI in the HIV−/METH− group (Dunnett’s posthoc test, p=0.019; Cohen’s d = −0.56), while there was no significant difference between the HIV−/METH− group and the other two groups (Fig 3B; Cohen’s d for HIV−/METH+ group = −0.22; Cohen’s d for HIV+/METH− group = −0.29). Furthermore, visual inspection of the data showed a stepwise decrease in PPI from the HIV−/METH− group to the HIV+/METH+ group, with the latter having the lowest PPI. Overall, these data suggest an additive effect of HIV and METH in reducing PPI in men.

The extremely low number of women in the HIV+/METH− group (n=5) and HIV+/METH+ group (n=2) would have produced statistical results with questionable value; however, we performed some statistical analyses and have included figures as supplementary material for the sake of completeness. We also included a table of demographic information for the women as supplementary material. Briefly, there was significant effect of prepulse intensity on PPI among women [F(2,82)=13.584, p<0.0005]. There was no prepulse intensity-by-HIV interaction, prepulse intensity-by-METH interaction, or prepulse intensity-by-HIV-by-METH interaction. As in the men, there was a main effect of HIV [F(1,41)=5.608, p=0.023], but no effect of METH or HIV-by-METH interaction.

### 3.9 Associations with HIV Illness Characteristics and Cognitive Performance

Among men, there were no significant correlations between average PPI and METH use characteristics or average PPI and HIV illness features. Average PPI in HIV+ men was negatively correlated with several cognitive domains, including learning, memory, motor function, and global deficit score (Table 2). These correlations were not corrected for multiple comparisons; however, these results are consistent with our predictions that worse PPI would be associated with worse cognitive function (note that higher scores on these cognitive tests indicate worse performance). No significant correlations were found between average PPI and cognitive domains in METH+ men. No significant correlations were found between average PPI and HIV illness features, average PPI and METH use characteristics, or average PPI and cognitive domains among women.

## 4. DISCUSSION

In this cross-species study of sensorimotor gating deficits as measured by PPI in HIV and METH, we found that METH treatment decreased PPI in male, but not female mice. PPI normalized with cessation of METH treatment. Furthermore, iTat transgenic mice exhibited normal PPI until they received DOX injections that induced Tat expression. At that point, male iTat mice exhibited decreased PPI relative to their littermates, but this effect was not seen in female iTat mice. There was no interactive effect of METH withdrawal and Tat expression on PPI in the iTat mice. In the human studies, HIV was associated with impaired PPI in both men and women. While the effects of METH on PPI were not statistically significant, we found evidence for an additive effect of HIV and METH on PPI, such that male HIV+/METH+ participants had the lowest PPI. We also found worse PPI was associated with worse performance in several cognitive domains among HIV+ men, but not the METH-dependent population.

Our results complement our previous study on the effects of HIV on sensorimotor gating in humans that found reduced PPI in HIV+ adults with HIV-associated neurodegenerative disorder (HAND), but not in the general HIV+ population (Minassian et al., 2013). It is possible the current study found impaired PPI in the general HIV+ population because the increased sample size afforded more power to detect the effect. Furthermore, the human and animal results of this study are in agreement in that HIV decreased PPI in men and iTat expression decreased PPI in male mice. These data suggest that the iTat mouse model at least partially replicates the effects of HIV on PPI in men and may be a worthwhile model to study gating deficits in HIV+ men. Alternatively, our animal results differ from those examining PPI in a different mouse model of HIV, the gp120 model. In this model, the HIV envelope protein gp120 is constitutively expressed in the brain from birth (Thaney et al., 2018). Henry et al. (2014) found that gp120 male mice exhibited similar PPI compared to WTs, but female gp120 mice had reduced PPI compared to WTs. Overall, these results show that PPI effects are specific to the HIV animal model in question, possibly due to the different HIV protein expressed in these models. Indeed, previous studies observed that intrahippocampal Tat injections reduced PPI in male but not female rats (Fitting et al., 2006), consistent with our current findings in mice. Alternatively, intrahippocampal injections of gp120 did not reduce PPI in rats (Fitting et al., 2007). Furthermore, intrahippocampal injections of Tat and gp120 have different effects on neuronal and glial cell number, with Tat, but not gp120, reducing neuronal number in the dentate gyrus, and Tat, but not gp120, increasing astrocyte number in multiple hippocampal regions (Fitting et al., 2008). Overall, our results indicate that HIV, and at least the induction of the Tat protein, is associated with decreased sensorimotor gating, with a possible influence of sex on this effect.

This study was to our knowledge the first to examine the effects of a history of METH dependence on PPI in humans. We did not see evidence of an effect. This observation was surprising given the cognitive deficits associated with METH dependence. It is worth noting that METH use characteristics (e.g. – days since last use, total quantity METH use) varied substantially between participants in this study, making it possible that a more homogenous group of people would show an effect of METH on PPI. That said, the human and animal data from this study generally agree with respect to the effects of METH on PPI. Most human participants in our study (~85%) had not used METH in seven or more days, suggesting that our population was most comparable to the mice during Phase 3, i.e. – after the cessation of METH use. Thus, both human and animal data indicate that, independent of HIV, there was no effect of METH on PPI after a period of METH cessation. The decreased PPI in male mice during METH treatment was consistent with previous studies whereby current METH treatment reduced PPI in male mice (Arai et al., 2008; Chao et al., 2012). However, other studies in mice observed that withdrawal from chronic METH treatment increased PPI (Henry et al., 2014). The discrepant results may have to do with the different METH treatment regimens, as these two studies used different patterns of METH injections. Henry et al. used an escalating, continuous (i.e. – daily) pattern of injections for 25 days with 7 days of withdrawal, while the current study used an escalating, but intermittent (i.e. – 4 days on, 3 days off) pattern of injections followed by 7 days of withdrawal. The interdose interval of methamphetamine treatment is known to affect behavioral outcomes (Kuribara, 1996). Therefore, the continuous versus intermittent treatment differences between these studies may account for the different results and require future direct comparisons.

Our study is also the first to examine the effects of combined HIV and history of METH dependence on PPI in humans. We found evidence that HIV and a history of METH dependence additively decreased PPI in men, and a similar trend was observed in women despite a very small sample size. Other studies have examined the effects of combined HIV and METH dependence on other CNS processes in humans and found mixed results. For example, clinical studies found that HIV and a history of METH dependence additively impaired cognitive function (Kesby et al., 2015; Rippeth et al., 2004). However, other clinical studies found that HIV and METH dependence did not additively impair everyday functional skills (Minassian et al., 2017) or impulsivity, disinhibition, or sensation-seeking (Marquine et al., 2014). Therefore, combined HIV and METH dependence likely has varying effects on CNS processes depending on the function examined. Overall, our current results suggest that comorbid HIV and a history of METH dependence additively impair early sensorimotor gating in humans. This deficit may contribute to impaired downstream cognitive processes, as seen in other studies. The animal studies, however, do not find an additive effect of HIV and METH. The animal results of the current study are consistent with a previous study whereby chronic METH did not worsen PPI in the gp120 mouse model of HIV (Henry et al., 2014). The different human and animal findings regarding additive HIV and METH effects may be due to the whole body infection with the full HIV virus seen in the humans versus expression of only one of the HIV proteins restricted to the brain in the animal models. Furthermore, there is evidence that chronic METH use may worsen consequences of HIV infection in humans in ways that animal models would not capture. For example, METH contributes to poorer adherence to ART (Reback et al., 2003) as well as biological ART resistance (Colfax et al., 2007). These factors may lead to greater neurotoxicity in humans with HIV compared to the gp120 or iTat models, and potentially account for the different results with respect to additive effects.

Finally, we examined associations between HIV illness characteristics, METH use features, cognitive function, and PPI. We found that worse PPI was associated with worse cognition. While these correlations were small and not corrected for multiple comparisons, the direction of the correlations was consistent with our hypothesis. Furthermore, the association of worse PPI with worse cognitive function was observed in a previous study population (Minassian et al., 2013). Finally, it was unexpected that there were no correlations between METH use characteristics and cognitive function, given that METH dependence is associated with impaired cognitive function (Potvin et al., 2018).

### 4.1 Strengths and limitations of this study

Strengths of this study include the extensive phenotyping of human subjects that allowed for detailed analyses of the relationship between HIV and METH dependence illness features and sensorimotor gating. Another strength of this study is its translational nature. By themselves, the human studies are only observational, limiting the ability to draw conclusions regarding causality. However, the animal experiments enable causal inferences to be made regarding the effects of HIV protein induction and METH use on sensorimotor gating. Also, the animal studies modeled METH escalation and binging, as well as an acquired HIV pathology, both of which are relevant to humans. Limitations of the study include the relatively few female subjects in the human component of the study, which is important since the mouse studies suggest sex effects that may interact with HIV and METH to influence sensorimotor gating. Furthermore, there was a lack of matching on age among the men in the human study. There was also substantial heterogeneity in the METH use patterns of subjects with a history of METH dependence. The animal model used also has limitations, as it only expresses a single HIV protein and is non-infectious.

### 4.2 Practical Implications

Our results suggest HIV and METH use additively impair sensorimotor gating in men, with evidence of a similar trend in women. This may have implications for cognition, since sensorimotor gating is thought to contribute to downstream cognitive function. Notably, the average time since last METH use in our human population was approximately five months, suggesting the additive effects of HIV and METH on PPI can persist long after acute intoxication. Therapies for targeting sensorimotor gating may be useful for improving cognitive deficits in the HIV+/METH+ population. Indeed, preclinical studies find that some agents, such as the atypical antipsychotic ziprasidone and the prolyl oligopeptidase inhibitor IPR19, improve deficits both in PPI and cognitive function (Abdul-Monim et al., 2003; Mansbach et al., 2001; Prades et al., 2017). It would be interesting to understand the effects of such compounds on the cognitive deficits associated with HIV and METH use. Overall, our results underscore that METH use in HIV+ individuals should be of concern to patients and physicians alike due to the additive effects on sensorimotor gating and potentially cognition.

### 4.3 Directions for future study

Future studies might investigate the effect of anti-retroviral therapy (ART) administration on PPI in humans or animals to determine if ART causally affects PPI. Furthermore, relatively little remains known about sensorimotor gating in people who actively use METH or who are acutely intoxicated. Administration studies in which METH is given to healthy controls, people with prior METH dependence, and/or people with HIV could provide further insight into how these conditions impact sensorimotor gating. Indeed, animal research suggests that METH intoxication is associated with decreased PPI, and there are no known human studies examining the effects of acute METH intoxication on PPI. Future animal studies could examine additional models of HIV, such as humanized mouse models (Marsden and Zack, 2017), to determine the effects of infection with the HIV virus versus isolated HIV proteins. Finally, given the sex differences found in the preclinical studies, future studies should examine the role of sex in the effects of HIV and/or METH on PPI.

### 4.4 Conclusions

This study determined the effects of chronic METH on PPI in the iTat mouse model of HIV and in people with HIV. Both chronic METH treatment and Tat expression reduced PPI in male, but not female mice. There was also no interaction between METH treatment and Tat expression in mice. We also found that HIV is associated with impaired sensorimotor gating in men and women. Furthermore, the combination of HIV and METH additively impaired sensorimotor gating in men. Finally, PPI was correlated with neuropsychological performance. These results suggest HIV+ patients who use METH are at risk of further impaired sensorimotor gating.

## ACKNOWLEDGMENTS

We would like to acknowledge Nathan Wood for his assistance with the human components of this study, as well as Mahalah R. Buell and Melissa Flesher.

## STATEMENT OF ETHICAL STUDY OF HUMAN SUBJECTS

This study was approved by the Institutional Review Board at the University of California San Diego. Subjects gave written informed consent for inclusion in the trial.

## SUPPLEMENTAL FIGURE AND TABLE LEGENDS

**Figure S1. Effects of HIV and a history of METH dependence on PPI in female participants.** Four groups of human women with or without HIV and with or without a history of METH dependence (HIV−/METH−: n=27, HIV−/METH+: n=11, HIV+/METH−: n=5, HIV+/METH+: n=2) were recruited and assessed for PPI at three different prepulse intensities. Average PPI for each group at each prepulse intensity are shown in panel A. PPI averaged across prepulse intensities are shown in panel B. Data are presented as mean ± S.E.M. * = main effect of HIV (ANOVA, p<0.05)

**Table S1: Demographic and Illness Features: Female Participants.** Data are presented as mean ± standard deviation and range (min-max) unless otherwise noted. ART: Anti-retroviral therapy, ARV: Anti-retrovirals; BDI-II: Beck’s Depression Inventory – 2^nd^ Edition; NA: Not applicable; NS: Not significant; WRAT: Wide Range Achievement Test

**Table S2: Demographic and Illness Features: All Participants.** Data are presented as mean ± standard deviation and range (min-max) unless otherwise noted. ART: Anti-retroviral therapy, ARV: Anti-retrovirals; BDI-II: Beck’s Depression Inventory – 2^nd^ Edition; NS: Not significant; WRAT: Wide Range Achievement Test

