## Supplementary figures and images for "Both HIV and Tat Expression Decrease Prepulse Inhibition with Further Impairment by Methamphetamine"

### Supplemental Fig 1

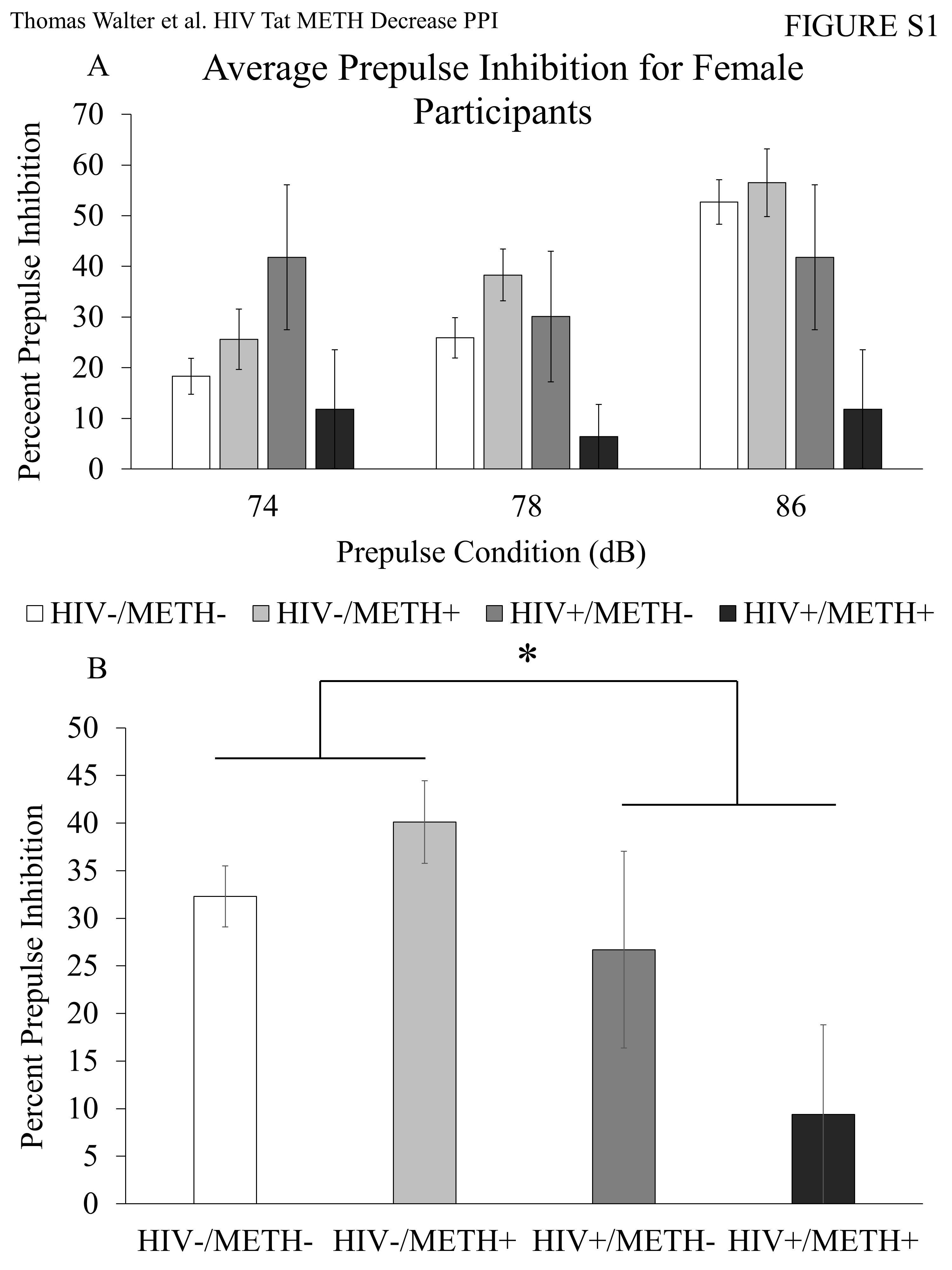
