## Supplemental Tables for "Both HIV and Tat Expression Decrease Prepulse Inhibition with Further Impairment by Methamphetamine"

Thomas Walter et al. HIV Tat METH Decrease PPI

**S1: Demographic and Illness Features: Female Participants**

|  | HIV-/METH-  (n=27) | HIV-/METH+  (n=11) | HIV+/METH-  (n=5) | HIV+/METH+  (n=2) | Statistics |
| --- | --- | --- | --- | --- | --- |
| Age (yrs) | 44.3 ± 18.5  (19-71) | 41.4 ± 11.3  (27-59) | 49.8 ± 8.6  (35-56) | 45.5 ± 26.2  (27-64) | NS |
| Ethnicity (n) | 4 Afr. Am.  6 Hisp.  2 Asian  14 Cauc.  1 Other | 1 Afr. Am.  5 Hisp.  0 Asian  4 Cauc.  1 Other | 0 Afr. Am.  0 Hisp.  0 Asian  4 Cauc.  1 Other | 0 Afr. Am.  1 Hisp.  0 Asian  1 Cauc.  0 Other | NS |
| Education (yrs) | 14.4 ± 21.7  (11-17) | 12.6 ± 3.0  (8-18) | 15.2 ± 2.2  (13-18) | 13.0 ± 1.4  (12-14) | **METH+ groups < METH- groups** |
| WRAT-4 Score | 106 ± 12  (87-126) | 93 ±11.5  (75-114) | 106 ± 12  (94-126) | 94 ± 1  (94-95) | **METH+ groups < METH- groups** |
| Global Deficit Score | 0.28 ± 0.27  (0-0.78) | 0.64 ± 0.52  (0.1-1.68) | 0.38 ± 0.40  (0.1-0.89) | 0.50 ± 0.55  (0.1-0.89) | NS |
| BDI-II Total Score | 4.9 ± 5.6  (0-18) | 13.4 ± 10.5  (0-29) | 16.4 ± 12.8  (2-35) | 13.0 ± 18.3  (0-26) | NS |
| Lifetime Substance Use Disorder (n) | 5 | 11 | 1 | 2 | **Chi-squared: p<0.0005** |
| Age First METH use (yrs) | - | 25.8 ± 11.0  (14-47) | - | 26.5 ± 16.2  (15-38) | NS |
| Days since last METH use (days) | - | 111 ± 136  (1-396) | - | 356 ± 118  (273-441) | **p=0.036** |
| Total days METH use (days) | - | 2706 ± 2433  (161-6886) | - | 5490 ± 4009  (2656-8325) | NS |
| Total quantity METH use (grams) | - | 3445 ± 5754  (117-19326) | - | 5330 ± 2712  (3411-7248) | NS |
| Current CD4+ Cell Count | - | - | 636 ± 400  (349-1285) | 796 ± 275  (602-991) | NS |
| Nadir CD4+ Cell Count | - | - | 246 ± 113  (90-356) | 390 ± 402  (106-675) | NS |
| Estimated Duration HIV Infection (yrs) | - | - | 8.9 ± 7.2  (2.5-16.7) | 4.5 ± 0.6  (4.1-4.9) | NS |
| Plasma Viral Load - log10(copies per mL) | - | - | 1.54 ± 0.91  (0.00-2.25) | 1.60 ± 0.00  (1.60-1.60) | NS |
| Current ART Use (n) | - | - | 5 Yes  0 No | 2 Yes  0 No | NA |
| Months Total ART Exposure | - | - | 73.4 ± 65.7  (11-180) | 54.3 ± 5.3  (50-58) | NS |
| Current Number of ARVs Using (n) | - | - | 3.2 ± 0.4  (3-4) | 3.0 ± 0.0  (3-3) | NS |
| Participants with AIDs (n) | - | - | 3 | 1 | NS |

Thomas Walter et al. HIV Tat METH Decrease PPI

**Table S2: Demographic and Illness Features: All Participants**

|  | HIV-/METH-  (n=69) | HIV-/METH+  (n=40) | HIV+/METH-  (n=52) | HIV+/METH+  (n=44) | Statistics |
| --- | --- | --- | --- | --- | --- |
| Age (yrs) | 39.1 ± 15.9  (18-71) | 38.2 ± 11.1  (21-60) | 41.8 ± 10.8  (23-70) | 39.7 ± 7.7  (24-64) | NS |
| Sex (n) | 27 F, 42 M | 11 F, 29 M | 5 F, 47 M | 2 F, 42 M | **Chi-squared: p<0.0005** |
| Ethnicity (n) | 10 Afr. Am.  16 Hisp.  2 Asian  39 Cauc.  2 Other | 9 Afr. Am.  10 Hisp.  0 Asian  20 Cauc.  1 Other | 9 Afr. Am.  13 Hisp.  2 Asian  26 Cauc.  2 Other | 4 Afr. Am.  11 Hisp.  0 Asian  28 Cauc.  1 Other | NS |
| Education (yrs) | 14.4 ± 2.0  (10-20) | 12.7 ± 2.4  (8-18) | 14.3 ± 2.1  (9-18) | 14.0 ± 2.6  (8-20) | **HIV-/METH+ < All other groups** |
| WRAT-4 Score | 105 ± 11  (82-131) | 96 ± 10  (75-116) | 103 ± 11  (83-130) | 105 ± 12  (87-131) | **HIV-/METH+ < All other groups** |
| Global Deficit Score | 0.23 ± 0.24  (0-0.95) | 0.44 ± 0.44  (0-1.68) | 0.34 ± 0.31  (0-1.11) | 0.48 ± 0.45  (0-1.67) | **METH+ groups < METH- groups** |
| BDI-II Total Score | 3.3 ± 4.5  (0-18) | 13.6 ± 10.6  (0-35) | 12.1 ± 13.2  (0-58) | 13.7 ± 12.1  (0-47) | **HIV-/METH- group < All other groups** |
| Lifetime Substance Use Disorder (n) | 25 | 40 | 23 | 42 | **Chi-squared: p<0.0005** |
| Age First METH use (yrs) | - | 23.3 ± 11.0  (8-56) | - | 25.1 ± 7.8  (13-48) | NS |
| Days since last METH use (days) | - | 173 ± 354  (1-2191) | - | 128 ± 135  (1-441) | NS |
| Total days METH use (days) | - | 2384 ± 207  (86-8797) | - | 1918 ± 204  (59-9771) | NS |
| Total quantity METH use (grams) | - | 3943 ± 7527  (35-40185) | - | 2117 ± 4386  (12-26526) | NS |
| Current CD4+ Cell Count | - | - | 644 ± 307  (81 – 1712) | 574 ± 235  (82 – 1053) | NS |
| Nadir CD4+ Cell Count | - | - | 312 ± 213  (3-800) | 287 ± 189  (8-840) | NS |
| Estimated Duration HIV Infection (yrs) | - | - | 10.3 ± 9.0  (0.1-28.9) | 8.2 ± 6.3  (0.1-23.8) | NS |
| Plasma Viral Load - log10(copies per mL) | - | - | 2.05 ± 1.19  (0.00-5.45) | 2.35 ± 1.38  (0.00-5.95) | NS |
| Current ART Use (n) | - | - | 40 Yes  12 No | 33 Yes  11 No | NS |
| Months Total ART Exposure | - | - | 70.8 ± 76.5  (0-275) | 59.0 ± 56.6  (0-181) | NS |
| Current Number of ARVs Using (n) | - | - | 2.7 ± 1.7  (0-6) | 2.7 ± 1.7  (0-5) | NS |
| Participants with AIDs (n) | - | - | 21 | 18 | NS |
